# Beyond the annotated: protein foundation models enable robust prediction of microbial root competence

**DOI:** 10.64898/2026.05.22.727091

**Authors:** Petra Matyskova, Gijs Selten, Corné M.J. Pieterse, Sanne Abeln, Ronnie de Jonge

## Abstract

**Background:** Root competence, the ability of soil bacteria to establish and grow on plant roots, is a key ecological trait influencing plant nutrition, growth, and health. However, identifying genomic determinants of root competence across bacteria remains challenging, in part because model generalisability depends strongly on how genomes are represented. Traditional approaches based on curated annotations are incomplete and biased toward well-characterised organisms and functions, limiting generalisation. Sequence-similarity clustering improves coverage but yields high-dimensional features relative to dataset size, hindering training. Foundation models offer an alternative by learning compact representations without relying on prior annotation.

**Results:** Here, we compared pretrained genome representations from protein and DNA foundation models (ESM-2, Bacformer, DNABERT-S) with annotation- and clustering-based features (KEGG orthology, OrthoFinder protein families) for predicting root competence using synthetic microbial community data from *Arabidopsis thaliana* and assessed generalisability across bacteria. When training and test sets contained taxonomically related bacteria, most approaches performed similarly. However, when test bacteria belonged to phyla entirely absent from training, reflecting high evolutionary separation across all levels of bacterial classification, only pretrained protein representations retained predictive performance. Bacformer-derived representations, which incorporate genomic context, supported the strongest generalisation, suggesting that conserved genomic organisation contributes to predicting root competence. Feature attribution quantifying protein contributions to model decisions linked root competence to TonB/SusD-dependent receptors, small-molecule transporters, and unannotated proteins with conserved regulatory motifs and homology to carbon starvation-response loci.

**Conclusions:** Protein foundation models support generalisation across evolutionarily distant bacteria and identify genomic determinants of root competence, including unannotated proteins.

## 1 Background

Plant diseases pose a significant challenge to crop yield and the agricultural economy [1], with fungal, bacterial, and viral pathogens affecting a wide range of crops. Current reliance on chemical pesticides for disease control has been shown to cause environmental harm, contribute to the emergence of resistant pathogens, and present risks to human health [2, 3]. Although genetic improvement and plant breeding have substantially improved crop productivity and disease resistance, plant health in agricultural systems is not determined by host genetics alone. Disease outcomes also depend on interactions with the surrounding microbial community, which can influence pathogen suppression, nutrient availability, and stress resilience [4]. The plant-associated microbiome therefore represents a complementary strategy improving plant performance and supporting sustainable agriculture [5]. Yet, the success of microbiome-based interventions inherently depends on a prerequisite process: root competence, the ability of soil microbiota to migrate to, establish on, and persist on the root [6].

Root competence is often studied within the context of complex microbial communities [7]. Although community assembly is influenced by stochastic processes such as dispersal limitation and priority effects, root microbiome communities repeatedly converge on similar compositions across identical host plants grown under similar environmental conditions [8, 9]. In parallel, root-associated microbial communities are consistently less diverse than the surrounding soil and represent a reduced subset of the broader soil microbiome [10]. This suggests that the plant root environment acts as a selective ecological filter shaped by host-associated processes and local physicochemical conditions [11]. While community context influences the final composition of the root microbiome, the capacity of individual microbes to pass this ecological filter is also determined by intrinsic physiological traits encoded in their genomes [6]. Understanding the genomic basis of root competence therefore represents a key step toward predicting and engineering beneficial plant–microbe interactions.

Early microbiome studies investigating microbial traits primarily relied on marker gene sequencing (e.g., *16S rRNA* profiling), which captures evolutionary relatedness but provides limited genome-level resolution [12, 13]. The increasing availability of microbial culture collections with whole-genome sequences and experimentally measured phenotypes enables genome-resolved analysis of microbial traits. Indeed, genomic features related to metabolism, motility, chemotaxis, and biofilm formation have been previously associated with root competence [14–16]. However, connecting microbial genomic content to ecologically relevant traits, such as root competence, while generalising across the evolutionary diversity of bacteria, remains challenging. Bacterial diversity is commonly described using hierarchical taxonomic ranks (e.g., phylum, class, order, family, genus), which group organisms by evolutionary relatedness [17]. A key factor shaping whether genomic signals generalise across these bacterial taxa is the way genome sequences are represented for analysis and modelling.

Genome sequences must be encoded into numerical representations before they can be used by predictive models. These representations determine which aspects of genomic data are accessible to the model, and therefore, shape both what patterns can be learned and how the resulting predictions can be biologically interpreted. Several strategies have been developed to represent microbial genomes. Functional annotation schemes (e.g., Gene Ontology (GO) [18], KEGG Orthology (KO) [19], PFAM [20], COG [21]) assign genes to known biological roles based on sequence similarity, producing interpretable and externally defined representations. However, these schemes vary in both specificity and coverage, with some annotations being too broad to be informative for microbiome research [22], and many genes remaining unannotated, particularly in poorly characterised organisms [23]. This limits the ability of predictive models to capture genome-wide signals and restricts biological insight into known gene–trait relationships. Furthermore, these representations have been shown to generalise poorly across the evolutionary diversity of bacteria [24]. In addition to many genes being unannotated, similar biological functions can be carried out by different gene families. As a result, the same biological process may be represented by different features, or not represented at all, in different bacterial taxa, making it difficult for models trained in one taxon to recognise equivalent functional signals in another.

Sequence similarity-based clustering approaches (e.g., OrthoFinder Groups (OGs) [25], OMA [26], Roary [27], Panaroo [28]) group genome-derived proteins into families without relying on prior functional annotations, achieving near-complete coverage and supporting post-hoc functional interpretation. However, these approaches produce sparse, high-dimensional representations relative to the number of genomes, which hinders the ability of predictive models to learn meaningful patterns. In addition, since gene families are inferred directly from sequence similarity within a dataset rather than from large, diverse reference databases, versions of the same gene may be split into separate clusters as sequences diverge across evolutionary groups [25, 29]. Consequently, the same biological function may be represented by different clusters in different bacterial taxa, leading to the same generalisation challenge observed for functional annotations. Taken together, traditional genome representation approaches struggle to simultaneously achieve manageable feature dimensionality, genome-wide coverage, and consistent representation across bacterial taxa.

Recent advances in protein and DNA foundation models offer a promising alternative by learning compact, information-rich representations directly from amino acid or nucleotide sequences, enabling the inclusion of unannotated genes. Protein foundation models (ESM-2 [30, 31], ProtTrans family models [32], Bacformer [33]) can capture functional, structural, and evolutionary signals in amino acid sequences. DNA foundation models (e.g., DNABERT family models [34], Nucleotide Transformer [35]) encode patterns related to genomic organisation, regulatory context, and nucleotide-level evolutionary signals, offering a complementary view. Representations derived from these models have demonstrated utility in tasks such as protein structure prediction [30], functional classification [36], and mutational effect estimation [37], suggesting their potential to generalise to trait-prediction applications. However, it remains unclear whether these pretrained representations capture signals that generalise across the evolutionary diversity of bacterial communities while remaining biologically interpretable.

In this study, we evaluate microbial genome representations for predicting root competence in *Arabidopsis thaliana*, using bacterial isolate genomes paired with experimentally-derived presence of bacteria in the rhizosphere. We assess functional annotation-based (KOs), clustering-based (OGs), and foundation model-derived representations with respect to predictive performance, biological interpretability, and generalisation across evolutionarily distant bacteria and cross-study variation. We evaluate ESM-2 as a general protein foundation model, Bacformer as a bacteria-specific protein foundation model that incorporates genomic context, and DNABERT-S as a species-aware DNA foundation model. We show that protein foundation model representations uniquely support robust predictive performance when models are tested on bacterial phyla absent from training, while effectively incorporating information from unannotated proteins and taking less computational time to derive. This establishes a framework for identifying genomic determinants of host–microbe interactions that may be missed by traditional representation schemes.

## 2 Results

### 2.1 Generalisation performance varies across evaluation settings

To compare microbial genome representations for predicting root competence, we fit the same multilayer perceptron architecture using the SynCom dataset described by Selten and colleagues [16]. This study quantified the ability of 988 individual bacterial isolates to colonise plant roots within controlled synthetic microbial communities, enabling comparative assessment of microbial recruitment under standardised conditions. The resulting measurements provide a consistent benchmark for evaluating genome-based predictors of root competence.

We compared functional annotation–based features derived from KEGG orthologues (KOs), sequence clustering-based features based on Orthofinder groups (OGs), and representations obtained from protein and DNA foundation models. Although annotation and clustering approaches produced sparse, high-dimensional feature spaces, foundation model representations achieved full genome coverage while being compact (320–768 features per genome) and substantially faster to extract (2–3 days on a single GPU) (Table 1). Here, we evaluated whether these representational differences translated into improved predictive performance across evaluation settings.

**Table 1.**
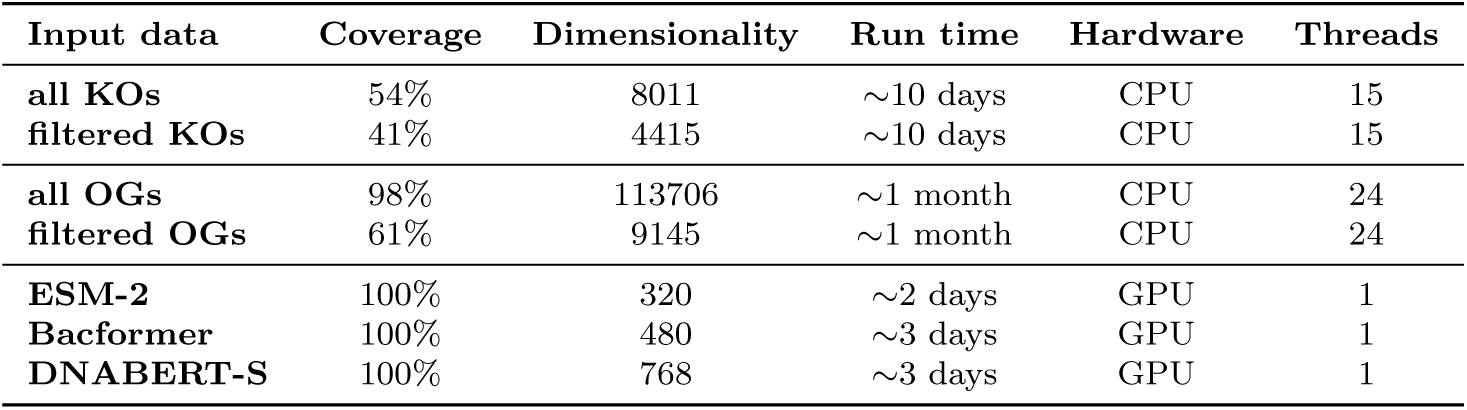
Comparison of microbial representation approaches based on coverage, dimensionality, and runtime. Summary of coverage (i.e., percentage of the sequence represented), dimensionality (i.e., the number of features per genome), and approximate computational runtime (i.e., the time required to generate representations for 988 bacterial isolate genomes). Coverage corresponds to the percentage of protein coding genes for the protein-based approaches and all genes for the DNA-based approach. Filters were applied to KOs (KEGG orthology groups) and OGs (Orthofinder protein families) to reduce the number of features, .05 meaning features with at least 5% variance across genomes are kept.

We evaluated model performance in three settings. We first tested performance on held-out bacteria taxonomically related to the training data, referred to as in-distribution (ID). Second, we evaluated performance on bacteria from phyla absent during training, referred to as out-of-distribution (OOD), representing generalisation across evolutionarily distant bacteria. Lastly, we also tested all models on six independent external SynCom datasets previously compiled and systematically analysed in Selten and de Jonge [38], representing generalisation across experimental contexts. An overview of the data, classification task, and evaluation settings is provided in Fig. 1.

**Fig. 1.**
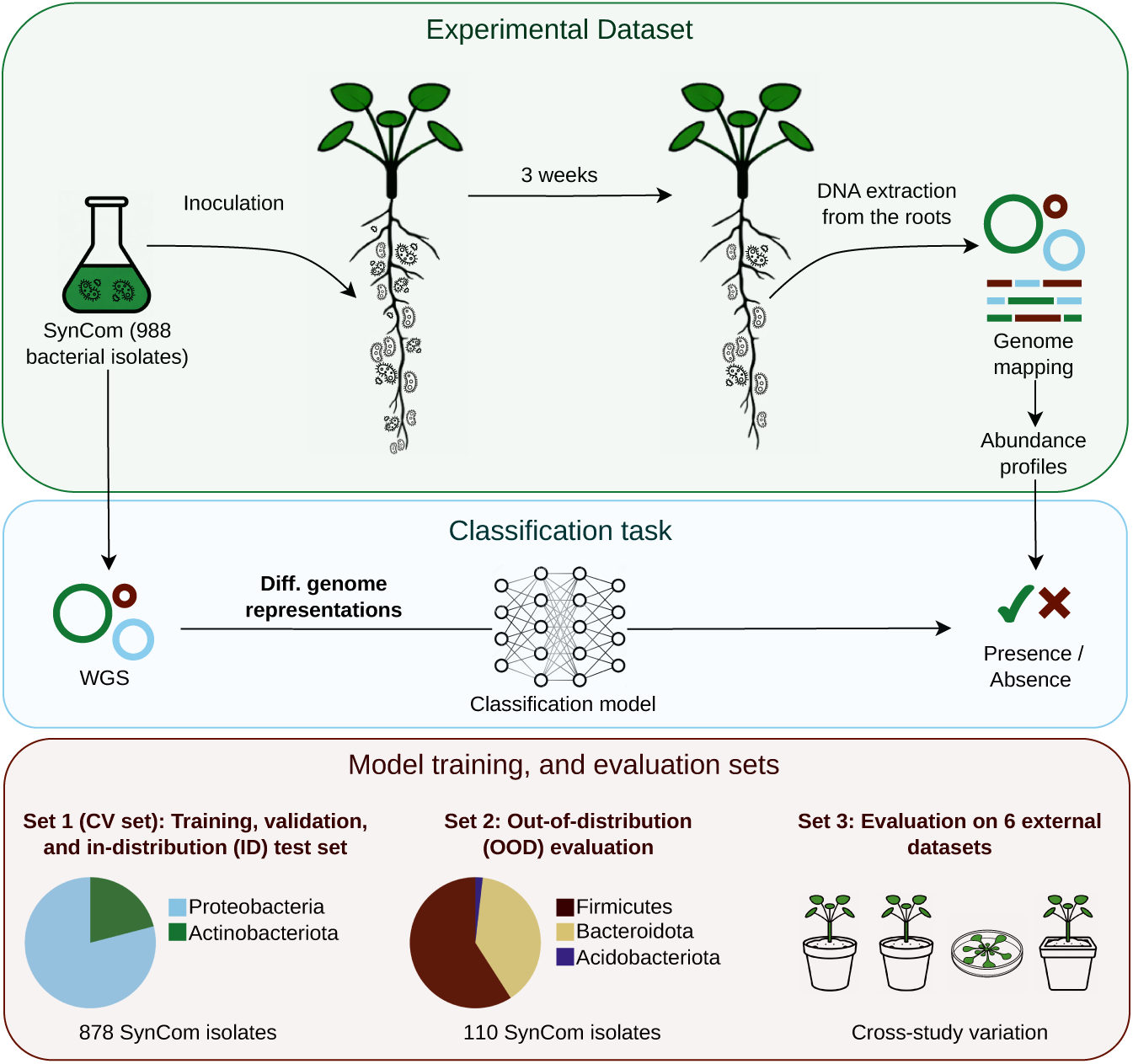
Overview of the experimental data, classification task, and model evaluation strategies. A synthetic community (SynCom) dataset from Selten et al. [16], linking bacterial iso-late genomes to experimentally-derived root colonisation ability in *Arabidopsis thaliana*, was used to train a root competence classifier from genomic information. Different whole genome sequencing (WGS)-derived representations were used as input to the same classifier architecture predicting bacterial presence or absence in the rhizosphere. Models were first trained and evaluated using cross-validation (CV, see Section 5.3) on Set 1, comprising of 878 out of 988 of the SynCom isolates (694 Proteobacteria, 184 Actinobacteriota), representing generalisation to taxonomically related bacteria. Performance was further assessed on Set 2, which consisted of the remaining 110 Syncom isolates from phyla absent in training (65 Firmicutes, 43 Bacteroidota, 2 Acidobacteriota), representing generalisation across taxonomically distinct bacteria. Set 3 corresponded to six independent external datasets (see Section 5.1.2), representing generalisation across experimental contexts. Pie charts represent the true taxonomic composition of the respective SynCom isolates.

#### 2.1.1 Models using functional annotations and foundation model representations perform comparably on evolutionarily related bacteria

Performance estimates on the ID test sets (Set 1, Fig. 1) revealed that models trained on KOs, Bacformer- and DNABERT-S-derived representations achieved similar performance (Table 2). Across these representations, the F1-scores ranged from 0.72–0.74 and the PR-AUC from 0.70–0.78. While individual metrics favoured different models, no single representation consistently outperformed the others across all performance measures.

**Table 2.** Performance on the ID (in-distribution) test sets. Performance of the same multilayer perceptron architecture on root competence task using different input data, KOs (KEGG Orthology) and OGs (Orthofinder protein families) being annotation- and clustering-based, and ESM-2-, Bacformer-, and DNABERT-S-derived representations being foundation model based. The best performance is represented in bold and second best underlined. Filters were applied to KOs and OGs to reduce the number of features, such that features with at least 5% variance across genomes are kept.

| Input data | Accuracy | Precision | Recall | F1 | AUC | PR AUC |
| --- | --- | --- | --- | --- | --- | --- |
| all KOs | **0.75 ± 0.02** | 0.70 ± 0.07 | **0.78 ± 0.04** | **0.74 ± 0.03** | **0.82 ± 0.03** | **0.78 ± 0.06** |
| filtered KOs | <u>0.74 ± 0.02</u> | 0.70 ± 0.06 | <u>0.75 ± 0.05</u> | <u>0.73 ± 0.03</u> | <u>0.81 ± 0.02</u> | **0.78 ± 0.04** |
| all OGs | 0.60 ± 0.01 | 0.58 ± 0.10 | 0.46 ± 0.03 | 0.51 ± 0.05 | 0.61 ± 0.05 | 0.60 ± 0.11 |
| filtered OGs | 0.58 ± 0.00 | 0.56 ± 0.09 | 0.46 ± 0.11 | 0.49 ± 0.05 | 0.62 ± 0.02 | 0.61 ± 0.09 |
| ESM-2 | 0.66 ± 0.03 | 0.61 ± 0.09 | 0.72 ± 0.06 | 0.66 ± 0.04 | 0.74 ± 0.01 | 0.69 ± 0.05 |
| Bacformer | <u>0.74 ± 0.04</u> | <u>0.72 ± 0.07</u> | <u>0.75 ± 0.08</u> | <u>0.73 ± 0.07</u> | 0.76 ± 0.06 | <u>0.70 ± 0.07</u> |
| DNABERT-S | **0.75 ± 0.00** | **0.75 ± 0.04** | 0.70 ± 0.06 | 0.72 ± 0.03 | <u>0.81 ± 0.02</u> | **0.78 ± 0.04** |

In contrast, OG–based models performed substantially worse (F1≈0.49–0.51; PR-AUC≈0.60–0.61), remaining close to random despite variance filtering. Models trained on ESM-2-derived representations showed intermediate performance (F1≈0.66; PR-AUC≈0.69), performing above OG-based models but below the rest (Table 2).

Together, these results establish a baseline: under favourable generalisation conditions within related bacteria, models trained on KO-based functional annotations and foundation model-derived representations capture comparable predictive signal, whereas OG-based representations fail to support accurate classification.

#### 2.1.2 Models using protein foundation model representations remain robust across evolutionarily distant bacteria

When evaluated on independent external SynCom datasets (Wolinska et al. [39], Finkel et al. (2020) [40], Hou et al. [41], Set 3 from Fig. 1), performance decreased relative to the ID evaluation but remained above random for models based on KOs and foundation model representations (Table 3). Across these datasets, KO-based models achieved F1-scores of 0.47–0.57 and PR-AUC of 0.64–0.69. In comparison, models based on filtered KOs generally showed reduced performance (F1: 0.27–0.35; PR-AUC: 0.59–0.63). Models using foundation model representations showed comparable performance ranges across datasets. ESM-2-derived representations led to F1-scores of 0.54–0.65 and PR-AUC of 0.67–0.72, while Bacformer-derived representations led to F1-scores of 0.58–0.63 and PR-AUC of 0.70–0.77. In comparison, DNABERT-S-derived representations supported similar F1 performance (0.54–0.65), although PR-AUC values were slightly lower (0.61–0.67). All models failed to generalise on three additional external datasets (Lebeis et al. [42], Finkel et al. (2019) [43], Duran et al. [44], Set 3 from Fig. 1), with all representations showing near-random predictive performance (Supplementary Table 4). While no single representation type consistently outperformed the others, models based on foundation model representations collectively achieved the highest or second-highest performance across metrics more frequently than KO-based models.

**Table 3.** Performance on external datasets and the taxonomically distinct OOD test set. Performance of the same multilayer perceptron architecture on root competence task using different input data, KOs (KEGG Orthology) being functional annotations and ESM-2-, Bacformer-, and DNABERT-S-derived representations being foundation model based. The OOD (out-of-distribution) test set consists of bacteria with no taxonomic overlap with the training set from phylum to genus. The best performance is represented in bold, second best underlined, and poor performance italic. Number of samples, n, in each dataset as well as number of negative samples and number of positive samples, neg./pos., is reported. Filters were applied to KOs to reduce the number of features, such that features with at least 5% variance across genomes are kept. The models failed to generalize on 3 out of 6 external datasets, Lebeis et al. [42], Finkel et al. (2019) [43], and Duran et al. [44] that were dropped from the table. All OG-based models also failed to generalise and were dropped for readability. The exact performance for all the omitted cases can be found in the Supplementary Table 4.

| Dataset | Input data | Accuracy | Precision | Recall | F1 | AUC | PR AUC |
| --- | --- | --- | --- | --- | --- | --- | --- |
| **Wolinska et al. [39]**<br>n=175, 80/95 | all KOs | 0.60 ± 0.05 | **0.68 ± 0.06** | 0.50 ± 0.08 | 0.57 ± 0.07 | 0.64 ± 0.05 | 0.64 ± 0.02 |
|  | filtered KOs | 0.49 ± 0.06 | 0.54 ± 0.08 | 0.26 ± 0.22 | 0.33 ± 0.20 | 0.59 ± 0.02 | 0.62 ± 0.04 |
|  | ESM-2 | **0.64 ± 0.04** | **0.68 ± 0.04** | <u>0.63 ± 0.06</u> | **0.65 ± 0.05** | <u>0.66 ± 0.03</u> | <u>0.67 ± 0.03</u> |
|  | Bacformer | 0.63 ± 0.01 | **0.68 ± 0.01** | 0.59 ± 0.01 | 0.63 ± 0.01 | **0.70 ± 0.01** | **0.71 ± 0.01** |
|  | DNABERT-S | 0.59 ± 0.01 | <u>0.61 ± 0.01</u> | **0.69 ± 0.05** | <u>0.64 ± 0.02</u> | 0.61 ± 0.02 | 0.64 ± 0.01 |
| **Finkel et al. (2020) [40]**<br>n=44, 23/21 | all KOs | 0.63 ± 0.07 | 0.70 ± 0.12 | 0.37 ± 0.15 | 0.47 ± 0.16 | 0.68 ± 0.06 | 0.69 ± 0.09 |
|  | filtered KOs | 0.53 ± 0.04 | 0.49 ± 0.08 | 0.21 ± 0.19 | 0.27 ± 0.19 | 0.61 ± 0.02 | 0.59 ± 0.04 |
|  | ESM-2 | 0.63 ± 0.04 | 0.70 ± 0.10 | 0.44 ± 0.07 | 0.54 ± 0.05 | 0.69 ± 0.02 | 0.72 ± 0.06 |
|  | Bacformer | **0.69 ± 0.01** | **0.83 ± 0.07** | 0.44 ± 0.03 | **0.58 ± 0.01** | **0.72 ± 0.01** | **0.77 ± 0.01** |
|  | DNABERT-S | 0.60 ± 0.08 | 0.61 ± 0.12 | **0.49 ± 0.06** | <u>0.54 ± 0.06</u> | 0.66 ± 0.05 | 0.67 ± 0.07 |
| **Hou et al. [41]**<br>n=175, 71/104 | all KOs | <u>0.57 ± 0.01</u> | **0.71 ± 0.03** | 0.47 ± 0.04 | 0.56 ± 0.02 | 0.60 ± 0.03 | 0.66 ± 0.04 |
|  | filtered KOs | 0.47 ± 0.06 | 0.62 ± 0.07 | 0.26 ± 0.19 | 0.35 ± 0.18 | 0.54 ± 0.05 | 0.63 ± 0.05 |
|  | ESM-2 | <u>0.57 ± 0.03</u> | 0.66 ± 0.03 | <u>0.56 ± 0.05</u> | <u>0.61 ± 0.03</u> | <u>0.62 ± 0.02</u> | <u>0.68 ± 0.03</u> |
|  | Bacformer | <u>0.57 ± 0.00</u> | 0.67 ± 0.00 | 0.54 ± 0.01 | 0.60 ± 0.00 | **0.65 ± 0.02** | **0.70 ± 0.02** |
|  | DNABERT-S | **0.58 ± 0.02** | 0.64 ± 0.00 | **0.67 ± 0.06** | **0.65 ± 0.03** | 0.58 ± 0.01 | 0.61 ± 0.01 |
| **OOD test set**<br>n=110, 88/22 | *all KOs* | *0.80 ± 0.00* | — | *0.00 ± 0.00* | *0.00 ± 0.00* | *0.50 ± 0.00* | *0.20 ± 0.00* |
|  | *filtered KOs* | *0.80 ± 0.00* | — | *0.00 ± 0.00* | *0.00 ± 0.00* | *0.50 ± 0.00* | *0.20 ± 0.00* |
|  | ESM-2 | 0.61 ± 0.31 | 0.40 ± 0.16 | 0.91 ± 0.16 | 0.53 ± 0.16 | 0.81 ± 0.10 | 0.54 ± 0.05 |
|  | Bacformer | **0.81 ± 0.02** | **0.52 ± 0.03** | **1.00 ± 0.00** | **0.68 ± 0.03** | **0.91 ± 0.01** | **0.62 ± 0.06** |
|  | DNABERT-S | *0.39 ± 0.05* | *0.05 ± 0.02* | *0.12 ± 0.07* | *0.07 ± 0.04* | *0.21 ± 0.02* | *0.13 ± 0.01* |

A contrasting pattern emerged on the OOD test set (Table 3, Set 2 from Fig. 1). Here, KO-based models exhibited poop performance, predicting the major-ity class (recall=0.00; F1=0.00; PR-AUC=0.20) exclusively, resulting in inflated accuracy (Acc=0.80) but no meaningful discrimination (ROC-AUC=0.50). Models based on DNABERT-S-derived representations similarly failed to retain predictive signal (F1=0.07 ± 0.04; PR-AUC=0.13 ± 0.01). In contrast, models using protein foundation model representations maintained substantial performance. Bacformer-derived representations supported the highest performance (F1≈0.68; PR-AUC≈0.62), while ESM-2-derived representations led to moderate predictive power (F1≈0.53; PR-AUC≈0.54).

OG-based models failed to achieve predictive performance across all external datasets and the OOD test set, collapsing to majority-class or near random predictions. These results are omitted from Table 3 and are reported in the Supplementary Table 4.

Together, these results show that while models based on KO functional annotations and foundation model representations can generalise comparably across cross-study variations, generalisation across evolutionarily distant bacteria is retained only by models based on protein foundation model representations.

### 2.2 Dataset similarity and foundation model representation analysis in the context of generalisation

The observed generalisation patterns beyond the training dataset raised two questions. First, we asked whether the preserved performance on three of the six external datasets reflects a smaller distribution shift with respect to the training dataset and whether a similar analysis could help explain the failure on the remaining datasets. Second, since models using representations derived from protein foundation models were the only ones to retain predictive performance across evolutionarily distant bacteria, we asked to what extent they capture taxonomic and orthogroup structure. To address these questions, we first quantified taxonomic overlap and representation similarity between the SynCom dataset used for training (Set 1, Fig. 1) and the other test sets (Set 2 and 3, Fig. 1). We then examined the extent to which protein representations recover taxonomic and protein family (orthogroup, OG) information.

#### 2.2.1 Taxonomic and representation-space similarity mirror predictive performance

To contextualise the generalisation patterns, we compared taxonomic and foundation model representation similarity of the test sets with respect to the training dataset (Fig. 2A). Genomes from the external datasets showed moderate taxonomic overlap with the genomes from training across ranks, with a proportion of shared taxa of 0.2–0.5 (Jaccard index). The highest proportion was at the order level, with a decrease both toward the highest (phylum) and lowest (genus) level. In contrast, genomes from the OOD test set exhibited no overlap across ranks, from phylum to genus, indicating no taxonomic overlap. This is consistent with how the OOD test set was constructed, excluding the least occurring taxonomies at the phylum level (see Section 5.1.1).

**Fig. 2.**
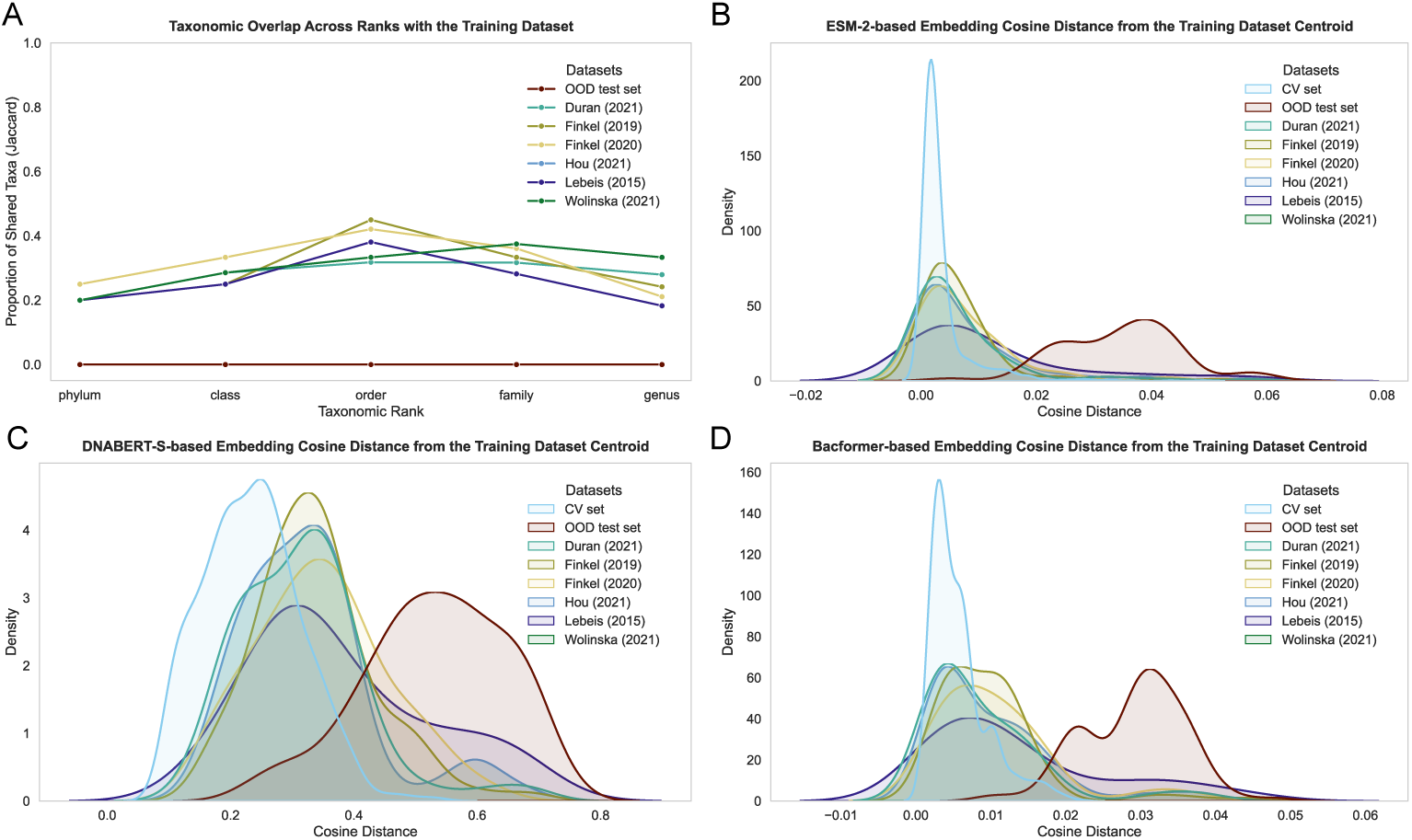
Dataset-level similarity between the training dataset and the other test sets. (A) Taxonomic overlap between the cross-validation (CV) dataset (Set 1, Fig. 1) and the other test sets (Set 2 and 3, Fig. 1), measured using the Jaccard index across taxonomic ranks. (B–D) Distribution of cosine distances between each genome representation and the cross-validation dataset centroid (mean representation) computed for genomes from all sets, using ESM-2- (B), DNABERT-S- (C), and Bacformer-derived representations (D).

Genome-level analysis of the foundation model representations (ESM-2, DNABERT-S, Bacformer) showed a similar trend. Representations of genomes from external datasets were close to the training dataset centroid (i.e., mean representation) in cosine distance, whereas the OOD test set displayed markedly larger distances (Fig. 2B-D). This structure mirrors the performance results, with external datasets representing a moderate domain shift and the OOD test set a large shift hindering generalisation.

#### 2.2.2 Protein foundation model representations recapitulate taxonomic and orthogroup structure

We assessed whether foundation model representations capture structured biological organisation at the genome and protein levels (Fig. 3). At the genome level, representations reflected hierarchical taxonomic organisation, with DNABERT-S-, ESM-2-, and Bacformer-derived representations capturing taxonomic structure to differing degrees (Fig. 3A). For ESM-2 and Bacformer representations, ARI and V-measure, which represent different measures of agreement between representation clusters and taxonomic labels (see Section 5.4.2), were relatively high at the phylum level, decreased at the class level, and progressively increased again toward order, family, and genus. DNABERT-S representations exhibited lower agreement at higher taxonomic ranks but increased toward the genus. Across ranks, Bacformer-derived representations generally achieved the highest scores, followed by DNABERT-S and ESM-2, with differences diminishing at lower taxonomic levels.

**Fig. 3.**
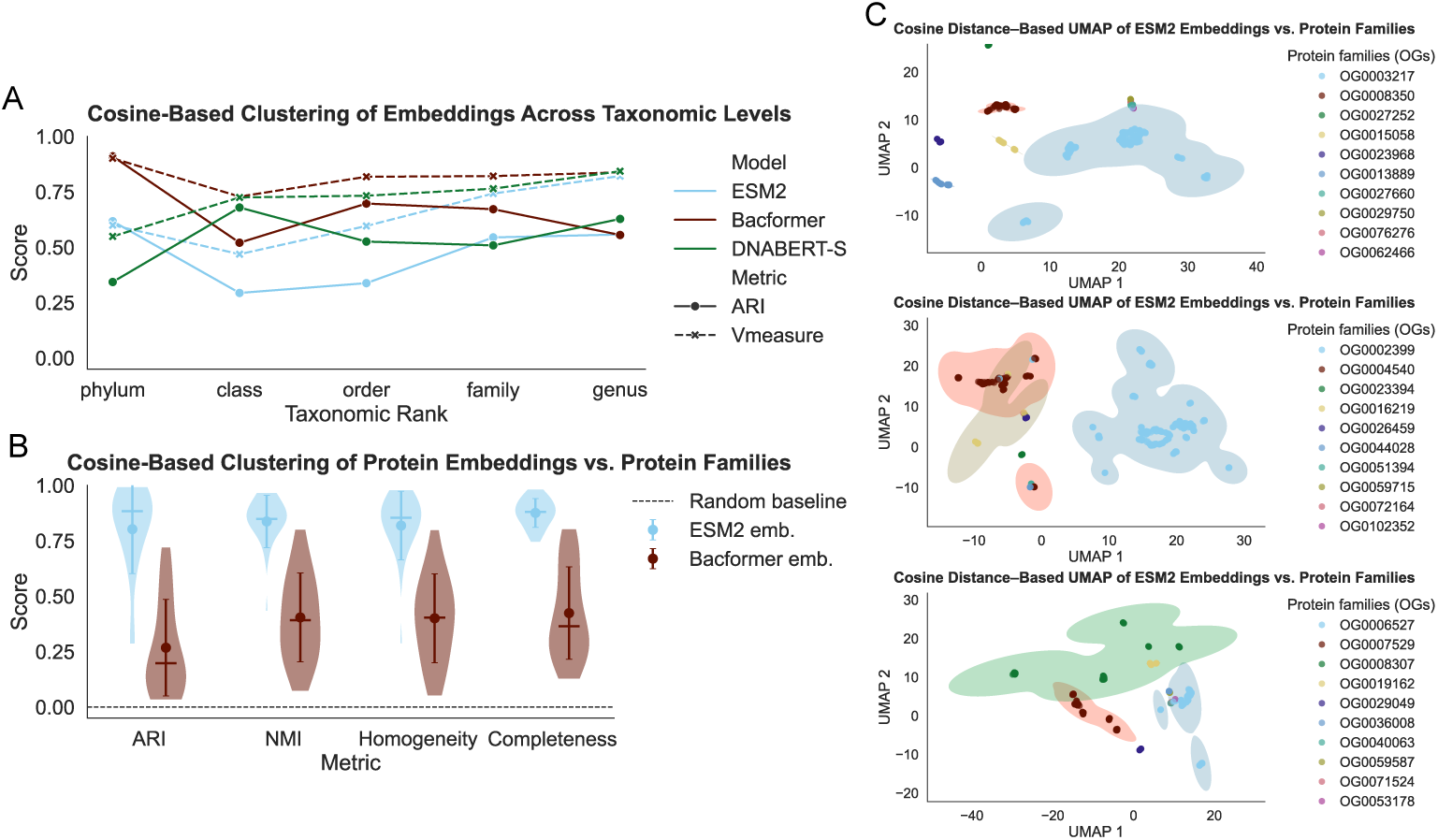
Orthogroup and taxonomic organisation captured by foundation model representations. (A) Genome-level cosine distance-based clustering performance of DNABERT-S-, ESM-2-, and Bacformer-derived representations/embeddings across taxonomic ranks, quantified using ARI and V-measure. For each model, genome embeddings were clustered using cosine distance, and resulting clusters were compared with known taxonomic labels at the phylum, class, order, family, and genus levels. (B) Protein-level clustering performance of ESM-2- and Bacformer-derived representations/embeddings compared to OG-based protein clustering. Genome embeddings were clustered using cosine distance, and resulting clusters were compared with OG-based protein families. Scores represent the distribution across 20 independent sub-sampling runs, each using proteins from 10 randomly selected OGs. The dashed line indicates a random baseline. (C) Cosine distance–based UMAP projections of ESM-2-derived representations/embeddings from three subsampling runs of B), illustrating the degree of cluster separation as compared to OG-based protein clusters.

At the protein level, representations showed clear correspondence with protein families (represented by OGs) across multiple clustering metrics (ARI, NMI, homogeneity, and completeness), which capture complementary aspects of agreement between computed clusters and defined protein groups, including cluster purity and the extent to which members of the same group are recovered together. The clustering of ESM-2-derived representations achieved the highest agreement, and the clustering of both protein foundation model representations consistently exceeded the random baseline (Fig. 3B). These results indicate that the representations recover orthogroup structure to a substantial extent. Cosine distance–based UMAP projections provide a qualitative visualisation of the representation space and show that proteins belonging to the same OG often occupy locally coherent regions of the projection (Fig. 3C), indicating that proteins from the same family tend to have similar representations. Since the projection compresses high-dimensional relationships into 2 dimensions, the visible separation between groups is only approximate and varies across protein families.

Together, these analyses show that representations derived from protein foundation models encode hierarchical taxonomic structure and orthology. However, these signals alone do not account for the observed differences in model generalisation. The partial but incomplete alignment with taxonomic ranks and OG-based protein families suggests that the representations capture additional continuous dimensions of biological variation beyond discrete taxonomic or orthogroup categories, which may facilitate the alignment of representations across larger taxonomic distances.

### 2.3 Model explainability identifies functional signatures associated with root-competent genomes

To identify genomic features underlying root competence, we performed an attribution-based explainability analysis using Integrated Gradients (IGs), which identify proteins that contribute the most to model predictions (see Section 5.5.1). We focus on the Bacformer-based model, which showed the strongest generalisation. The analysis distinguished between class-specific signals, used by the model when predicting each phenotype, and differential signals, separating root-competent from non-root-competent genomes. Biological interpretation focused on the most stringent subset of discriminatory signals (level 2), meaning protein groups that not only showed the strongest contribution to model predictions (level 1), but were also recurrently and significantly overrepresented among the model’s top contributors in one class relative to the other (FDR ≤ 0.05, see Equation 9 for formal criteria).

#### 2.3.1 Shared metabolic cores and rhizosphere-specific functional biases

We examined how the classifier internally recognises each phenotype by analysing attribution patterns separately for root-competent and non-root-competent genomes, thereby capturing class-specific signals independent of their discriminatory value.

Across both phenotypes, high-attribution pathways mainly reflected core metabolic and growth-associated functions (Fig. 4A in gray). Glycolysis/gluconeogenesis and the pentose phosphate pathway supported central carbon processing and redox balance, while oxidative phosphorylation, sulfur metabolism, and porphyrin-linked cofactor biosynthesis indicated energy conservation and respiratory capacity. Shared investment in biosynthetic machinery was reflected by nucleotide (purine/pyrimidine), fatty acid, and ribosomal pathways. These findings characterise the genomes as metabolically competent and respiration-capable, with strong biosynthetic capacity.

**Fig. 4.**
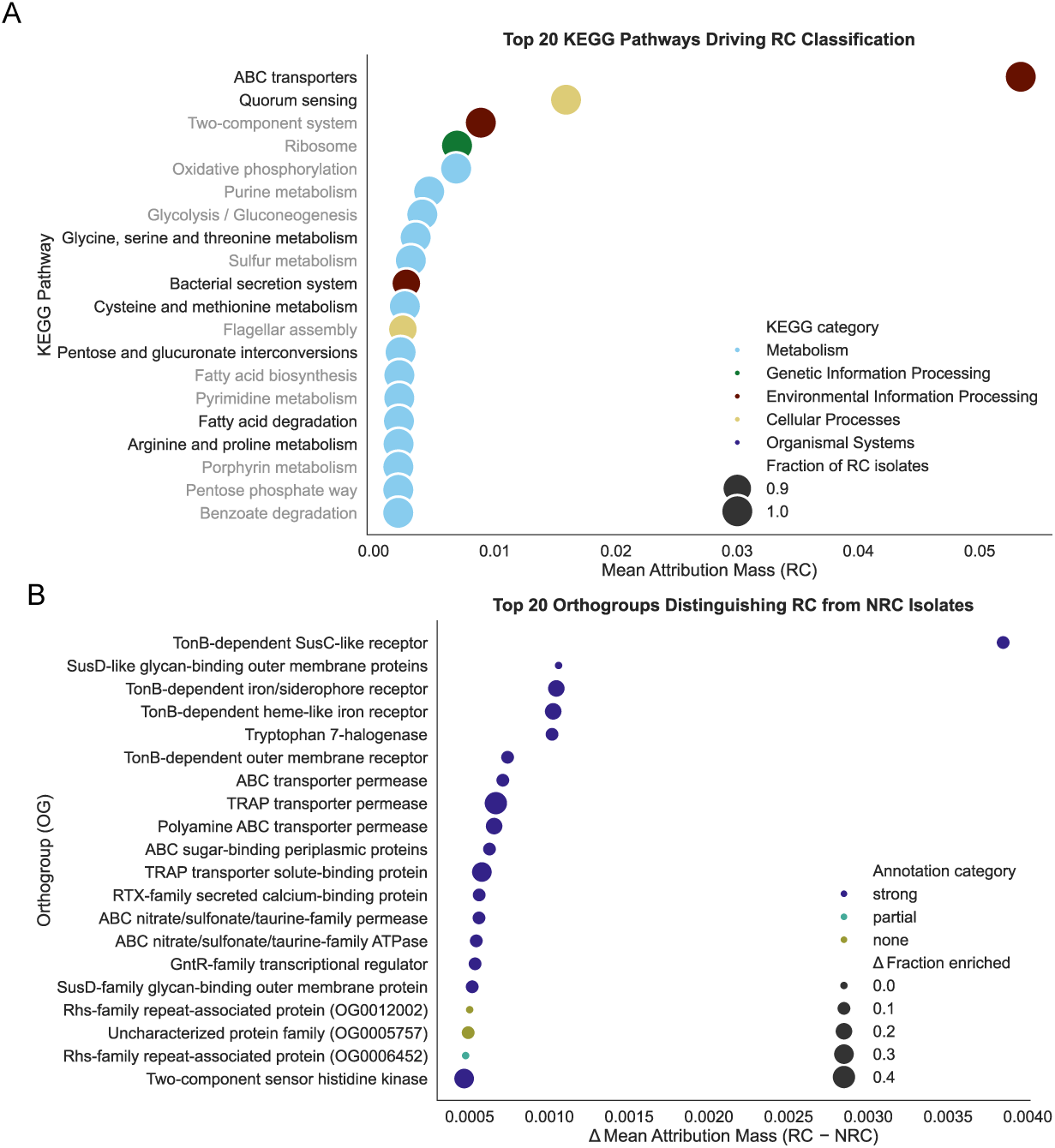
Model-derived functional signals of root-competent genomes across KEGG pathways and OGs. Explainability analysis of the best generalising model trained on Bacformer representations. Protein contributions to model predictions were quantified using Integrated Gradients (IGs). Proteins in the top 50% of IG scores were grouped into KEGG pathways or OGs (Orthofinder protein families). For each group, we computed global mean attribution mass (Equation 5) and prevalence (fraction of genomes containing ≥1 retained protein). A) Top 20 KEGG pathways associated with root competence (RC) classification, ordered by mean attribution mass, consistent with class-specific analysis criteria (Section 5.5.1). Pathways in gray were also driving the non-root competence (NRC) classification (Supplementary Fig. 7). B) Top 20 OGs distinguishing RC from NRC genomes, ordered by the difference in mean attribution mass (Δ RC–NRC), consistent with the Level 2 differential analysis criteria (Equation 9). OGs are coloured by annotation category: strong (≥3 KO annotated proteins), partial (≥5 or *<*3 KO annotated proteins), and none (*<*5 annotated proteins). Group names denote consensus functional labels from available annotations.

The root-competent phenotype showed high-attribution pathways related to amino acid interconversion (glycine, serine, threonine, arginine, and proline), sulfur and one-carbon metabolism (cysteine/methionine), and diversified carbon utilization (pentose/glucuronate and fatty acids) (Fig. 4A). These signatures indicate increased metabolic flexibility, supporting substrate switching and redox modulation. Additionally, enriched pathways for ABC transporters, quorum sensing, and secretion systems suggest enhanced nutrient uptake and environmental interaction (Fig. 4A). Collectively, this profile reflects adaptation to nutrient-diverse, competitive environments where metabolic plasticity and regulatory responsiveness confer a selective advantage.

In contrast, non-root-competent genomes were characterized by biosynthetic self-sufficiency and centralised energy metabolism, including expanded amino acid biosynthesis, folate production, and TCA cycle activity, suggesting adaptation to resource-limited environments (see Supplementary Fig. 7A for full results). Overall, the class-specific attribution profiles were biologically coherent, reflecting conserved, high-abundance protein families that serve as stable within-class anchors. These pathways provide the model with reliable structural information for class prediction rather than phenotype-specific ecological signals.

#### 2.3.2 High-confidence markers distinguish rhizosphere-associated signatures

To identify the specific protein families that selectively differentiate root competence, we performed differential analysis with high-stringency (level 2) criteria at the OG level. This granularity allows the identification of marker proteins independently of curated annotation status (see Supplementary Fig. 7 for KEGG pathway-level results).

The top root-competent OGs align with a plant-root specialist strategy (Fig. 4B). The most prominent signal comprises a nutrient-import module for macromolecules, specifically SusC/SusD-like glycan-binding proteins and TonB-dependent receptors for glycans, hemes, and iron/siderophores. A second root-competent module features systems for low-molecular-weight solute uptake, including TRAP-family components and diverse ABC transporters for sugars, polyamines, and organosulfonates. These proteins facilitate metabolic adjustment to fluctuating small-molecule availability, supported by GntR-family and sensor histidine kinase regulatory factors. Finally, root-competent genomes encode an interaction-associated module containing RTX-family and Rhs-repeat secreted proteins alongside tryptophan 7-halogenase. Collectively, these features describe a genomic capacity for flexible substrate acquisition, dynamic regulation, and competitive interaction; consistent with adaptation to the rhizosphere environment.

A notable signal associated with root competence was OG0005757, an uncharacterized orthogroup that emerged as a top predictor despite lacking functional annotation (Fig. 4B). Although BLASTp top hits returned hypothetical proteins, lower-ranking alignments revealed conserved regulatory motifs including helix–turn–helix, PDDEXK, α/β barrel, and tRNA-synthetase–like domains, suggesting nucleic acid–interacting or regulatory functions. PaperBLAST [45] linked this group to the CCNA 00822/CC 0781 loci in *Caulobacter crescentus*, regulated by the carbon starvation mediator CspC [46]. This suggests that rapid metabolic state transitions may contribute to root competence.

In contrast, the top non-root-competent OGs reflect a strategy toward persistence and nutrient conservation. This profile includes dormancy modules, nonspecific adhesion and structural modules, and systems for the uptake of diffusible substrates (see Supplementary Fig. 7B for full results). Together, differential attribution indicates that the model prioritizes plausible mechanistic signals to distinguish between phenotypes. These findings highlight genomic differences associated with root competence and the model’s ability to identify discriminatory features linked to divergent ecological strategies.

## 3 Discussion

Identifying the genomic traits of root competence is essential for microbiome-based agriculture, as it enables the selection and engineering of plant health and growth-promoting beneficial microbes that can reliably colonise plant roots and enhance crop performance under diverse environmental conditions. However, prediction from genomic sequence alone faces several challenges. First, traditional representation schemes often have incomplete coverage and/or high dimensionality, which can hinder model learning [23, 24]. Second, these representations may fail to recognise equivalent features across bacterial taxa, hindering generalisation. Additionally, standard evaluation within taxonomically similar bacteria or on random test sets risks that associations merely reflect evolutionary relatedness rather than true genomic determinants of root competence [24, 47–49]. To address these limitations, we evaluated whether the dense genome-wide coverage of foundation model-derived representations can support generalisation beyond training data while remaining biologically interpretable.

More specifically, we compared functional annotation- and clustering-based features with representations derived from DNA and protein foundation models for predicting microbial root competence in *Arabidopsis thaliana* SynCom experiments. When taxonomically similar bacteria were present in both training and testing, KOs (KEGG orthology groups), Bacformer, and DNABERT-S supported broadly comparable predictive performance, whereas OGs (Orthofinder protein families) led to poor, and ESM-2 to intermediate performance. However, across taxonomically distinct bacteria, when test bacteria belonged to phyla absent from training, only models using protein foundation model representations retained predictive performance. This suggests that at least some genomic determinants of root competence extend beyond broad taxonomic structure, including unannotated genes.

Across all representations, models failed to generalise on three of six external datasets for reasons not evident from taxonomic or representation space analysis, possibly reflecting experimental, ecological, or growth-condition variation. The comparable performance of models which used KOs and foundation model representations on the taxonomically similar test sets should be interpreted in light of an important asymmetry: foundation model representations were derived without task-specific fine-tuning, and therefore represent a baseline rather than an upper bound on predictive capacity [50]. OG-based models performed poorly even on the taxonomically similar test sets despite full proteome coverage, possibly due to extreme feature dimensionality and sparsity. However, since variance filtering reduced the OG dimensionality to a KO-comparable range without recovering performance, dimensionality alone cannot explain the difference. Since these models already trained poorly, their failed generalisation should be interpreted in that light. Meaning, it is difficult to determine whether the collapse on external and OOD test sets reflects failed generalisation due to learning dataset-specific signals rather than transferable biological determinants, or a broader inability of the models to learn effectively from OG-based representations. The weaker performance of models based on ESM-2-derived representations relative to Bacformer across evaluation settings may reflect generalist training of ESM-2 on UniRef sequences biased toward eukaryotes and the absence of genome-context information [30]. Bacformer addresses both through bacterial-specific retraining and the incorporation of local genomic neighbourhood information, although its advantage may partially reflect the larger underlying ESM-2 base model [33]. The failure of KO-based models to generalise across bacterial taxa matches previous findings but may also reflect the susceptibility of high-dimensional sparse binary representations to learning broad co-occurrence patterns, including taxonomically structured ones, rather than finer task-specific signals. Finally, the generalisation failure of models based on DNABERT-S may reflect DNA-level representations predominantly capturing compositional and structural genomic signatures rather than functional information [34], previously found to be a key predictor of root microbiome assembly [16].

Having established that models based on protein foundation model representations retain robust predictive performance, we interrogated the best-performing Bacformer-based model to identify determinants of root competence. An important strength of this study is that root competence was assessed in a complex SynCom context, allowing the identified genomic signals to be interpreted as traits relevant to community-associated root competence. Attribution analysis distinguished between signals used by the model within each phenotype and differential signals separating the phenotypes. Although orthogroup-based representations performed poorly as standalone predictive features, orthogroup clustering remained useful for interpretation, grouping evolutionarily related proteins into biologically coherent units beyond curated annotation space. Accordingly, OGs were used alongside KEGG-based annotations to support the interpretation of attribution signals at the protein family level. In the class-specific analysis, both phenotypes showed high attribution for core bacterial competencies such as central carbon metabolism, respiration, and strong biosynthetic capacity, alongside additional functional patterns that differed; root-competent genomes exhibited greater emphasis on metabolic flexibility and environmental interaction, and non-root-competent genomes showed biosynthetic self-sufficiency and centralised energy metabolism. These within-class profiles are consistent with known adaptation mechanisms [51, 52], validating that the model relies on meaningful signals.

At the most stringent differential level, the analysis identified high-confidence genomic determinants of root competence. Consistent with the other results, phenotype separation was not driven by the presence or absence of major functional categories but by distinct implementations of them, likely involving different protein families, transport systems, and regulatory modules. Both phenotypes invest in surface attachment, nutrient acquisition, and adaptation to resource limitation, but differ in the ecological character of these investments. Non-root-competent genomes were characterised by dormancy machinery, nonspecific adhesion and structural modules, and regulatory systems oriented toward diffusible low-molecular-weight substrates, consistent with persistence under chronic nutrient limitation with greater reliance on diffusible or community-modified resources [51, 52]. Root-competent genomes exhibited diverse TonB/SusD-dependent uptake of complex substrates, flexible small-molecule transport systems, rapid-response regulatory systems, and interaction-associated proteins, consistent with adaptation to the temporally pulsed and chemically heterogeneous substrate availability of the rhizosphere and the competitive interactions it sustains [51–53]. A notable finding was the prioritisation of OG0005757, an uncharacterised orthogroup, among the strongest root-competence-associated predictors. Deeper BLASTp alignments suggested a regulatory protein architecture, and homology-based literature mining linked this group to CspC-regulated carbon starvation–response loci in *Caulobacter crescentus*, consistent with the rhythmic nutrient dynamics of the rhizosphere [54–56]. The emergence of an unannotated protein family as a high-confidence predictor highlights the discovery potential of approaches not restricted to annotated protein space.

Several limitations should be considered when interpreting these findings. Root competence was defined as a binary classification of a continuous competence phenotype in a single host species under controlled conditions, discarding quantitative information and limiting inference to other hosts, root morphologies, soil types, or field settings. All isolates were derived from root-associated communities; therefore, non-root competence should be interpreted as reduced competence on *A. thaliana* under tested conditions rather than strict soil adaptation. While bulk soil communities could serve as an alternative negative control, their dynamic and condition-dependent composition introduces interpretive ambiguities absent in the current design, as organisms in transit to or from roots or with untested competence potential coexist with established soil residents [57]. Attribution-based explainability analysis was restricted to correctly classified genomes and identifies model-supported associations rather than causal mechanisms, requiring experimental validation. Genome-level representations do not capture microbe–microbe interactions that shape community assembly [58], and should therefore be interpreted as reflecting individual genomic capacity rather than community-level outcomes. Representations were derived by mean pooling, which discards genomic organisation and regulatory context; however, more advanced aggregation approaches remain constrained by variable protein content and limited data. The analysis is restricted to bacteria, leaving other microbial kingdoms unaddressed.

Despite these limitations, the ability of protein foundation model-based representations to recover biologically coherent and experimentally testable signals suggests that this framework may help prioritise candidate determinants of microbiome-associated traits for future mechanistic and gene perturbation studies.

## 4 Conclusion

In summary, our results show that protein foundation model-derived representations provide compact and robust features for microbial trait prediction, outperforming annotation-based, clustering-based, and DNA-level approaches in their ability to generalise across evolutionarily distant bacteria while enabling the recovery of signatures that extend to unannotated proteins. Although root competence was evaluated in a single host system, the explainability analysis revealed broad ecological tendencies and specific genomic determinants of bacterial phenotypes. This work highlights the potential of sequence-based foundation models to advance genomic prediction in settings with limited annotation coverage, high taxonomic diversity, and modest dataset size, where dense, low-dimensional representations are particularly effective and traditional approaches struggle.

## 5 Methods

### 5.1 SynComs: Synthetic Community Datasets

#### 5.1.1 Training dataset

To train and evaluate the models predicting root competence based on representations derived from microbial genomes, we used a synthetic community (SynCom) dataset previously generated and described by Selten et al. [16]. The dataset comprises 988 bacterial isolates whose genomes were sequenced and subsequently cultivated together. This SynCom (referred to as SSC in the original paper) was inoculated onto *Arabidopsis thaliana* plants, enabling experimental quantification of each isolate’s abundance in the root compartment after three weeks of growth. The abundance measurements, combined with the corresponding isolate genomes, establish a direct link between genomic content and root competence.

The 988 bacterial isolates originated from three host-specific culture collections, *Arabidopsis thaliana* (AtSC), *Hordeum vulgare* (HvSC), and *Lotus japonicus* (LjSC), previously assembled from healthy plants grown in natural soils at Reijerscamp (NL), Askov (DK), and Cologne (DE), respectively. All isolates were whole-genome sequenced prior to assembly into the SynCom inoculum. Plants were cultivated under controlled growth-chamber conditions using a sterile sand/gravel/vermiculite substrate. Following inoculation, roots were harvested, DNA was extracted, and metagenomic sequencing was performed by mapping metagenomic reads to the isolate genome database using Salmon [59] and normalised by genome length as described previously [16]. Although the SynCom was inoculated onto *A. thaliana*, *H. vulgare*, and *L. japonicus*, only the *A. thaliana* subset was used in this study. The resulting dataset therefore contains paired information for each isolate: its genome sequence and its experimentally determined relative abundance in *A. thaliana* roots.

Normalised isolate abundances were subsequently converted into binary presence–absence labels to facilitate classification of root competence. Two thresholds were evaluated (0 and 0.0001), and a threshold of 0 was retained as it produced more balanced class distributions and higher model performance. For biological replicates, each replicate was binarised individually, and the majority label across replicates was used as the final class assignment for each isolate.

To assess model generalisability across evolutionarily distant bacteria, we constructed a taxonomically disjoint split of the dataset. A set of 110 genomes, also referred to as the OOD (out-of-distribution) test set, was identified based on having no taxonomic overlap with the remaining genomes from phylum to genus. The selection was made from the least occurring phyla that would together lead to an approximate 90/10 to 80/20 train/test split. The 110 genomes comprising 65 Firmicutes, 43 Bacteroidota, and 2 Acidobacteriota were fully excluded from all stages of model training. The remaining 878 genomes comprising 694 Proteobacteria and 184 Actinobacteriota were used for the nested cross-validation training procedure. Within each outer fold, performance was evaluated on the corresponding held-out subset of these 878 genomes, referred to as the ID (in-distribution) test set. This design enabled systematic estimation of within-lineage predictive performance while providing an independent evaluation on lineages absent from training. Further details on model architecture, feature construction, and the nested training scheme are provided in Section 5.3.

#### 5.1.2 External datasets

To assess model generalisability beyond the training dataset, we evaluated performance on six publicly available SynCom studies, Duran et al. [44], Finkel et al. [40, 43], Hou et al. [41], Wolinska et al. [39], and Lebeis et al. [42]. These studies represent diverse experimental designs, growth substrates, and host conditions, while providing both isolate genomes and community composition data. The selected datasets focus on *A. thaliana* and collectively capture SynComs cultivated on calcined clay (Duran et al., Lebeis et al.), FlowPot systems (Hou et al., Wolinska et al.), and agar media (Finkel et al.).

For each study, the isolate genomes and the corresponding community sequencing data were retrieved as described by Selten and de Jonge [38], who standardised isolate quantification using the SyFi pipeline [60] and normalised read counts by *16S rRNA* copy number. Only *A. thaliana* samples representing the control condition in each study were kept for evaluation, ensuring comparability with the training dataset, which reflects microbial competence under untreated, baseline conditions. The same binarisation procedure described for the training dataset was applied, with replicate samples treated individually prior to majority voting. In cases where replicate measurements showed two distinct abundance states, the majority state was retained to avoid diluting consistent outcomes across replicates.

### 5.2 Microbial representations

To compare alternative microbial representation approaches, we evaluated traditional annotation- and clustering-based features with foundation model–derived representations. The process of constructing representations and model training is illustrated in Fig. 5.

**Fig. 5.**
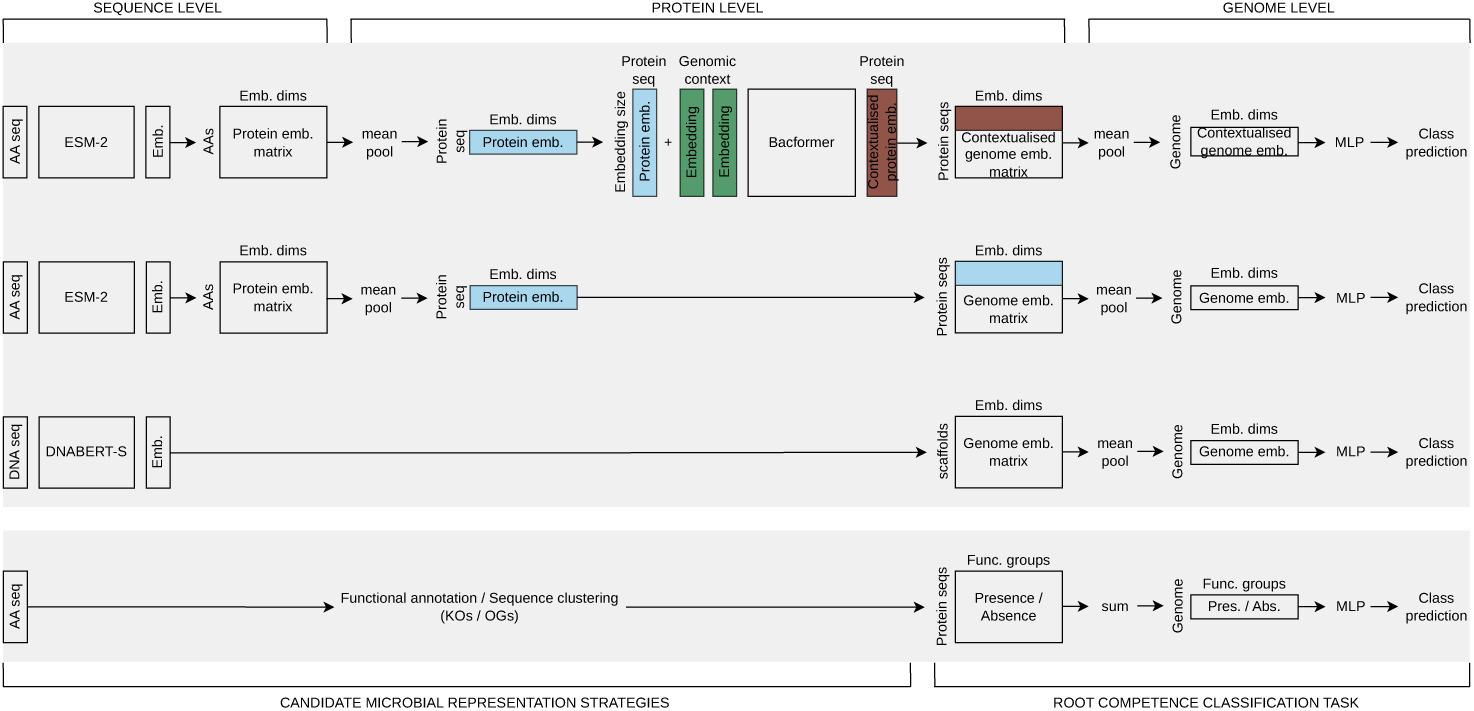
Sequence-to-classification pipelines for the foundation model-derived compared to annotation- and clustering-based approaches. Pipelines for models based on foundation model representations on top, with Bacformer, ESM-2, and DNABERT-S, top to bottom; and traditional approaches at the bottom. For Bacformer representations, amino acid sequences are encoded with ESM-2 and mean-pooled to protein-level vectors, concatenated with genomic-context representations and processed by Bacformer specific transformer layers yielding contextualised protein vectors. These are stacked into a genome-level matrix and mean-pooled to obtain a genome-level representation vector. For ESM-2 representations, per–amino acid ESM-2 vectors are stacked and mean-pooled to protein-level vectors; then stacked into a genome representation matrix, and mean-pooled to a genome-level representation vector. For DNABERT-S representations, DNA sequences are encoded using DNABERT-S to produce scaffold-level vectors, stacked into a genome-level matrix and mean-pooled to a genome-level representation vector. For representations obtained using traditional approaches, PROKKA-derived protein sequences were aggregated into families to obtain KOs (EggNOG annotation-derived KEGG orthology groups) and OGs (Orthofinder protein families). Protein-family presence/absence variation was summed to a genome-level representation vector. In all cases, the final genome-level representation vector provides the input to the same multilayer perceptron architecture to predict root competence.

Traditional representations were obtained from protein sequences derived from genome assemblies using Prokka [61]. The proteins were then grouped into families using two complementary pipelines: 1) eggNOG-mapper [62], which assigns KOs (KEGG Orthology groups), covering roughly half of all proteins (Table 1), among other functional categories; and 2) OrthoFinder [25], which clusters proteins into OGs (orthologous protein families) based on sequence similarity, resulting in near complete coverage but much higher feature dimensionality (Table 1). Both representations were binarised to indicate the presence or absence of each family per genome. To reduce sparsity, an additional filtered variant was tested by retaining only features with at least 5% variance across genomes (Table 1).

For foundation model representations, DNA sequences and PROKKA-derived protein sequences were encoded using pretrained transformer models. At the protein level, we used ESM-2 [30], trained on UniRef50, and Bacformer [33], a bacterial-specific ESM-2-based model informed by genomic neighbourhood context. At the DNA level, we used DNABERT-S [34], which creates species-informed representations of genomic scaffolds, capturing sequence organisation and regulatory context. For the protein models, foundation model outputs were first mean-pooled across amino acids to create per-protein representations. For all models, the representations were then mean-pooled across all proteins or scaffolds, yielding one vector representation per genome used as model input (Fig. 5). Representation vectors were inspected for outlier dimensions, and filtered vs. unfiltered versions were evaluated; the best performance was consistently obtained using all feature dimensions.

Traditional and foundation model-derived representations were also compared in terms of coverage, dimensionality, and approximate runtime (Table 1). Coverage refers to the proportion of proteins successfully represented; dimensionality to the number of genome-level features; and runtime to the approximate time required to generate all representations, without specific optimisation for computational efficiency.

### 5.3 Classification model

All microbial representations were evaluated using the same feed-forward multilayer perceptron architecture to ensure that performance differences reflected varying representations rather than model capacity. The network comprised three fully connected layers with LeakyReLU activation and dropout, followed by a single output neuron and sigmoid activation with a standard 0.5 threshold producing genome-level root competence predictions. Model weights were initialised using Kaiming normal for hidden layers and Xavier normal for the output layer. Models were trained using binary cross-entropy loss and optimised with either Adam, for foundation model representations, or stochastic gradient descent (SGD), for traditional representations. Preliminary experiments using a unified optimizer across representation types did not yield stable convergence. Specifically, SGD failed to converge on dense foundation model-derived inputs under a broad range of learning rates, while Adam failed to achieve stable training on high-dimensional sparse KO- or OG-based profiles. We therefore selected the optimizer that consistently achieved convergence within the nested cross-validation framework. Early stopping and adaptive learning-rate scheduling were applied to prevent overfitting.

Model optimisation followed a nested cross-validation scheme (Supplementary Fig. 8) on the training dataset comprising 878 genomes (Set 1 in Fig. 1). The outer cross-validation loop (three folds) estimated generalisation performance, while the inner loop (three folds) tuned hyperparameters using Optuna [63]. For each outer fold, three inner Optuna runs were performed, one per inner fold, to identify the best-performing hyperparameters based on validation loss. The selected configuration was then used to retrain a final model on the combined training and validation data, which was subsequently evaluated on a held-out test partition. Early stopping, learning-rate scheduling, and Optuna’s pruning strategy were employed to stabilise training and limit overfitting. Hyperparameters explored included dropout rate and weight decay; and L1 regularisation strength and batch normalisation on the first layer. Full model parameters and hyperparameter information are listed in the Supplementary Hyperparameters.

Model performance was assessed on each outer-fold held-out test set using standard classification metrics: Accuracy, Precision, Recall, F1-score, ROC-AUC, and PR-AUC. Precision, Recall, and F1 were calculated for the positive class (root-competent genomes). Reporting multiple complementary metrics ensured robustness to potential class imbalance and comparability across datasets.

### 5.4 Analysis of dataset structure and foundation model-derived representations

#### 5.4.1 Dataset-level similarity analysis

To explore potential causes of generalisation differences between datasets, we compared the cross-validation dataset (Set 1, Fig. 1) with the other test sets (Set 2 and 3, Fig. 1) in terms of both bacterial taxonomic and representation similarity.

Taxonomic similarity between datasets was assessed across multiple ranks (phylum, class, order, family, and genus). To quantify compositional similarity between datasets, we calculated Jaccard index, defined as the proportion of taxa shared between two datasets relative to the total number of unique taxa present in either. Jaccard index was computed on presence–absence data, treating each taxon as either present or absent in a given dataset. For interpretability, the fraction of taxa shared with the training dataset at each taxonomic rank was used, representing the proportion of its taxonomic diversity recovered in other datasets.

To complement the taxonomy-based analysis, we quantified how far each test set lies from the cross-validation dataset distribution in the foundation model representation space. We defined a reference centroid as the mean representation of all genomes in the training dataset and computed, for every genome in each dataset (Sets 1 to 3, Fig. 1), the cosine distance between its representation and this centroid. The resulting per-genome cosine distances were summarized as one-dimensional kernel density estimates stratified by dataset.

#### 5.4.2 Representation-level analysis

To evaluate the biological structure encoded in the foundation model-derived representations, we assessed their correspondence to OG-defined protein groups and to established taxonomic classifications. First, we examined how closely representation-derived clusters align with established taxonomic ranks. Pairwise cosine distances between genome embeddings were computed to generate a genome–genome distance matrix, which was subjected to agglomerative clustering. The resulting cluster labels were compared to known taxonomy using ARI and V measure across phylum, class, order, family, and genus. ARI measures overall clustering agreement [64] while V-measure summarises cluster purity and completeness with respect to the reference taxonomy [65]. This analysis provides a quantitative measure of the degree to which foundation model representations recover taxonomic structure at different evolutionary depths.

Second, we measured how well representation-derived clusters reproduce known OG assignments. Because the full annotation set spans more than 100,000 OGs, we used a repeated sub-sampling strategy: in each of 20 independent runs, 10 OGs were selected at random, and all proteins belonging to these OGs were included. Pairwise cosine distances were computed between protein representations within each run, and the resulting distance matrix was used for unsupervised agglomerative clustering. Cluster assignments were compared to previously obtained OG labels using Adjusted Rand Index (ARI), Normalized Mutual Information (NMI), homogeneity, and completeness. Metrics were aggregated across runs to obtain a robust estimate of the orthogroup structure captured by the representations.

### 5.5 Explainability analysis

#### 5.5.1 Model explainability via Integrated Gradients

To quantify the contribution of individual proteins to genome-level predictions, we computed Integrated Gradients (IGs) [66] for the best-performing model. The whole explainability pipeline is illustrated in Figure 6. IGs were computed individually for all correctly classified genomes with respect to the model’s output probability for the root-competent class, using the mean representation of all non-root-competent genomes as a biologically meaningful baseline. Since the model operates on an input matrix of N proteins by M feature dimensions, raw IG attributions were obtained for each protein at every dimension. IG was selected over SHAP because the classifier’s input is a genome-level representation derived via linear mean-pooling of individual protein representation. For such linear aggregations, IG and SHAP converge to identical values within the framework of additive feature attribution. The additive property of Shapley values ensures that the genome-level attribution is distributed linearly among its constituent proteins, making SHAP mathematically equivalent to gradient-based methods in this architecture. Consequently, SHAP would provide no additional explanatory power while incurring an exponentially higher computational cost to resolve attributions across thousands of proteins. IG thus provides a theoretically principled, additive framework that remains computationally tractable, maintaining similar axiomatic guarantees as SHAP [67].

**Fig. 6.**
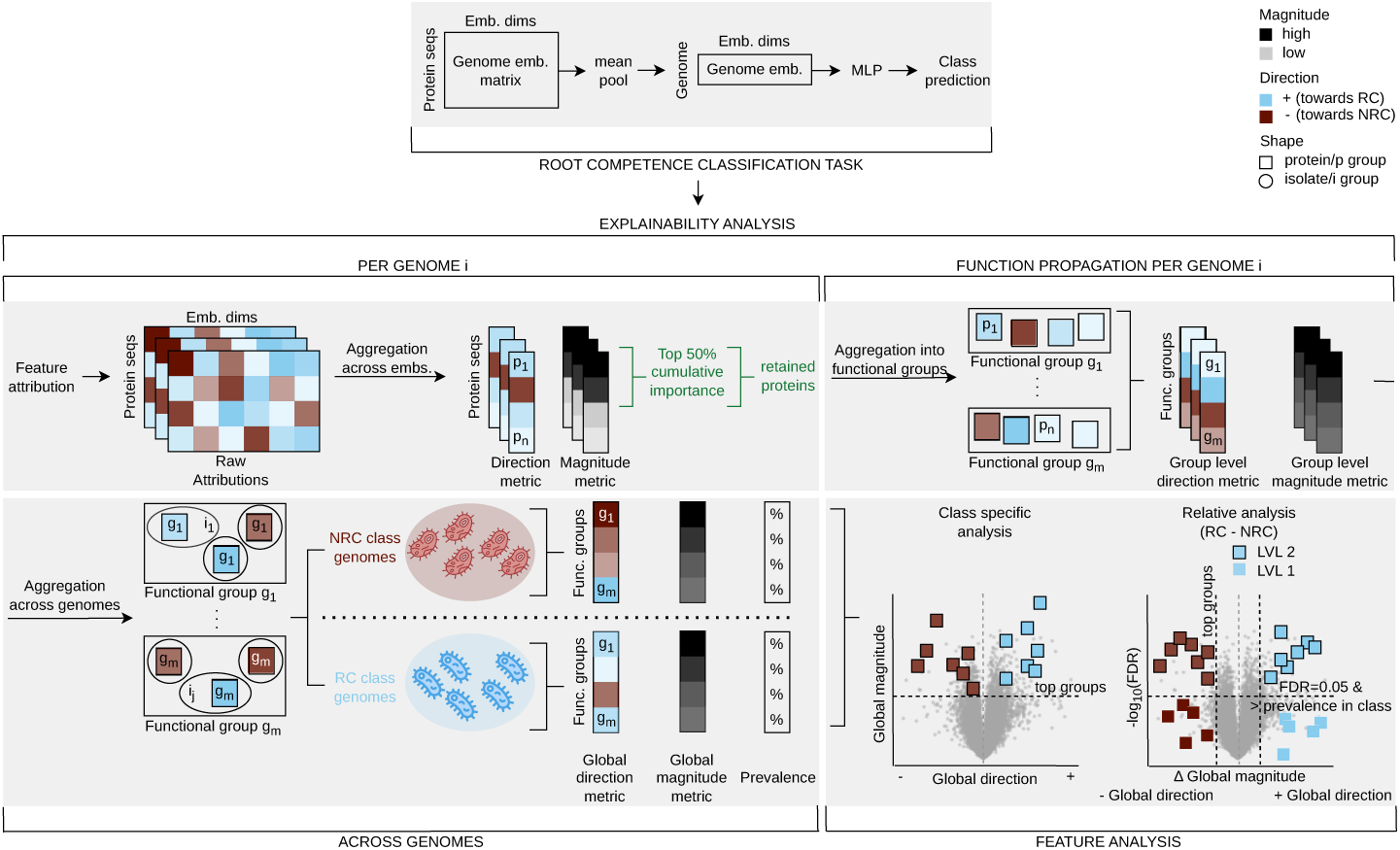
Overview of the Integrated Gradients–based explainability framework for genome-level protein attribution. The pipeline illustrates how IGs (integrated gradients) are computed and aggregated to interpret model decisions. For each correctly classified genome, IG attributions are computed with respect to the root-competent (RC) class using the mean protein representation across non-root-competent (NRC) genomes as baseline, producing per protein attribution vector. (Middle left) Raw IG attribution values are aggregated across feature dimensions to obtain a direction metric (Signed IG, Section 5.5) and a magnitude metric (Mean Absolute IG, Section 5.5). Proteins are ranked by magnitude, and the minimal set explaining the top 50% of cumulative normalized attribution is retained (Eq. 1). (Middle right) Protein families from traditional approaches are propagated to retained proteins, enabling computation of group-level direction (Eq. 2) and group-level magnitude metrics (Eq. 3) within each genome. (Bottom left) These values are then aggregated across genomes within each class to obtain global direction metric (Eq. 4), global magnitude metric (Eq. 5), and prevalence (Section 5.5) of each protein group among the retained proteins. (Bottom right) Two complementary analyses are performed. Class-specific analysis characterises the model’s internal logic within each phenotype independently and is based on the global magnitude metric requiring class directional consistency criteria (Section 5.5). Relative analysis identifies class differential features (RC vs. NRC) and is performed at two levels. Level 1 is based on differential global magnitude metric (RC-NRC) (Eg. 7) requiring class directional consistency criteria (Eq. 8). Level 2 analysis is additionally based on differential prevalence (Eq. 6) also requiring the prevalence related FDR ≤ 0.05 (Eq. 9).

From the resulting raw IG attributions, two complementary per protein (*p*) metrics were derived: 1) Signed IG (SignedIG), obtained by summing the IG attributions across feature dimensions, indicating the direction of influence (positive towards the root-competent class, negative towards the non-root-competent class); 2) Mean Absolute IG (MeanAbsIG) obtained by averaging the absolute value of the IG attributions across feature dimensions, representing the magnitude of influence irrespective of direction, providing a stable estimate when positive and negative contributions partially cancel.

To enable comparability across genomes with different protein counts and attribution scales, we normalized attribution magnitudes within each genome. Specifically, for each bacterial isolate *i* with protein set *P_i_* we computed a normalized absolute attribution per protein *p* ∈ *P_i_* as

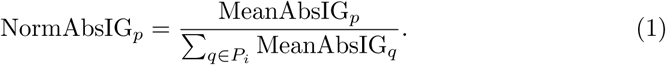

This ensures that 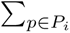 NormAbsIG*_p_* = 1 for every genome. Proteins were then ranked in descending order by NormAbsIG and a cumulative sum (CumNormAbsIG) was computed. To focus the analysis on the dominant contributors to the genome-level prediction task, we retained proteins until CumNormAbsIG ≤ 0.5, i.e., the minimal set of proteins explaining approximately the top 50% of the cumulative normalized absolute attribution within that genome.

Additionally, protein family assignments (KEGG pathways and OGs) were propagated to the retained proteins. Within each genome, we computed group-level metrics for each protein family as follows. Let *P_i,g_* denote the set of retained proteins in bacterial isolate *i* that belong to protein group *g*. 1) Mean signed IG defined as

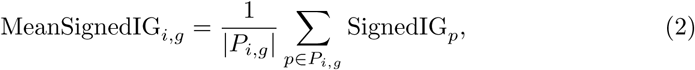

which quantifies the average directional effect of a protein group per genome, indicating whether its retained proteins consistently push the prediction toward root competence or non-root competence. 2) Attribution mass defined as

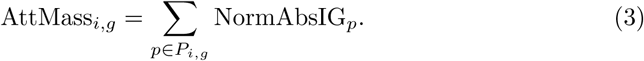

Since the normalized absolute attribution sums to one per genome, the attribution mass measures the fraction of the genome’s total attribution accounted for by the group’s retained proteins. This reflects how much of the model’s core reliance is placed on this protein group within the top-attribution set.

To analyse the model behaviour globally, we aggregated genome-level metrics by protein groups across genomes. For each class *C* ∈ {RC, NRC}, we defined *I_g,C_* as the set of correctly classified bacterial isolates in which 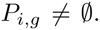. Class-level summaries were then derived by averaging metrics over *I_g,C_* as follows: 1) Global mean signed IG attribution defined as

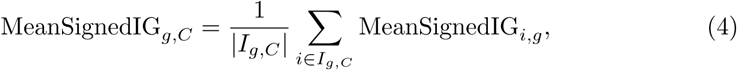

which captures whether the proteins belonging to a group consistently drive predictions toward the root-competent or non-root-competent phenotype across genomes. 2) Global mean IG attribution mass defined as

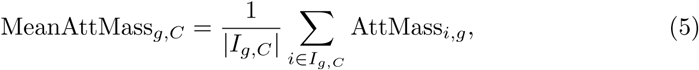

which reflects the average importance concentrated in a group’s retained proteins. This represents the average proportion of total attribution a group accounts for, specifically when its members are among the primary (top 50%) predictors. 3) Prevalence, defined as the fraction of correctly classified genomes of class *C*, in which at least one retained protein belonged to group *g*, capturing how frequently a group’s proteins are retained as top contributors across genomes.

To interpret grouped attributions and separate within-class model focus from between-class discriminatory signals, we defined two complementary analyses: 1) Class-specific analysis, quantifying the internal logic of the classifier within each class independently. 2) Relative analysis, identifying discriminatory features by comparing the root-competent and the non-root-competent class. For the relative analysis, we computed differences in group prevalence as

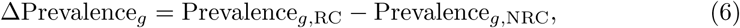

and the global mean attribution mass as

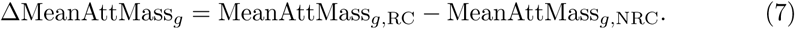

Additionally, to test whether a protein group occurs among the top-attribution proteins significantly more frequently in one class than the other, we performed a two-sided Fisher’s exact test. For each protein group, we compared the counts of correctly classified bacterial isolates in each class where 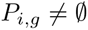 against the total number of correctly classified bacterial isolates in each class. Resulting p-values were corrected for multiple testing using the Benjamini–Hochberg procedure (FDR ≤ 0.05).

Analysis was restricted to data meeting specific quality and consistency criteria. First, only correctly classified genomes were included to ensure that IG attributions reflected successful model decision-making. Second, protein groups were required to satisfy class-consistency criteria. For class-specific analysis, we required a positive global mean signed IG for groups in root-competent genomes and negative global mean signed IG for groups in non-root-competent genomes, ensuring that the proteins from the group consistently pushed toward the predicted class. This constraint prevents the inclusion of groups with inconsistent or noisy patterns.

Relative analysis was conducted at two levels of granularity to identify model-supported class determinants. Level 1 identifies protein groups (*g_Cl_*_1_) characterized by differential attribution mass. A group was assigned to this level if it was more heavily weighted and directionally consistent for its associated class:

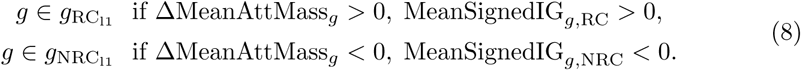

The results of this analysis can be found in Supplementary Fig. 7. Level 2 identifies the most stringent discriminatory markers (*g_Cl_*_2_) by requiring consistency across four metrics, including statistical enrichment:

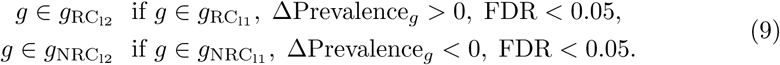

These stringent criteria ensured that downstream biological interpretations reflect robust, model-supported determinants where the group is more frequently used, more heavily weighted, and directionally consistent for its associated class.

#### 5.5.2 Model explainability across protein annotation quality

To interpret the model’s reliance on specific proteins, we propagated the protein family assignment described in Section 5.2 to the retained proteins. We utilized two systems to balance interpretability and precision: KEGG pathways provide broadly understood metabolic contexts, while OGs offer higher resolution by capturing orthologous groups, including those without known functions.

In the KEGG pathway-level analysis, we manually curated several labels that are marked with a latin cross sign. KEGG pathway names are known to reflect historical research bias toward human, plant, and animal systems, such that conserved prokaryotic enzymes are often assigned to eukaryote-specific pathway maps. This can result in apparent mismatches. To prevent misinterpretation, we manually annotated these maps with generalized, mechanism-centric names that describe the KO-supported molecular functions rather than the original organism-specific labels. This renaming does not alter the KO-level mapping and serves only to correct annotation bias and improve interpretability in a bacterial context.

For OG-level analysis, we quantified the annotation status of each OG to characterise the model’s decision-making across well-studied and uncharacterised protein space. Each OG was categorised based on the functional information of its member proteins, with strong annotation (containing KOs), partial annotation (containing GO terms, EC numbers, or PFAM domains but no KOs), or no annotation. An OG was labeled as strongly annotated if it contained at least three proteins with strong annotation. A partial annotation was assigned if the OG contained fewer than three strongly annotated proteins but at least five partially annotated proteins. All other OGs were categorised as unannotated.

OGs were assigned group-level functional annotation label based on functional information of the constituent proteins. For strongly annotated OGs, KO labels were aggregated to derive a consensus label. For partially annotated OGs, all available protein-level annotations were consolidated to generate a representative group-level label. OGs classified as unannotated but containing at least one annotated member were also assigned a group-level label based on the available functional information.

To further interpret unannotated protein space and illustrate how foundation models facilitate biological discovery, we selected OG0005757 from the top 20 discriminatory root competence features, which lacked annotations across all its proteins, and implemented a tiered functional annotation strategy. For a subset of the proteins, we first applied DeepFRI [68], a structure-based protein function prediction, to obtain Gene Ontology (GO) term predictions. However, the resulting annotations were either absent or limited to highly generic GO terms that did not provide meaningful functional insight. We next queried InterPro [69] to identify conserved domains or family signatures, which also yielded no informative matches. To further explore potential structural homology, AlphaFold [70] predicted structures were generated and searched against structural databases using Foldseek [71]; however, no informative structural analogs were identified. Finally, to assess homology more broadly, BLASTp searches against NCBI’s non-redundant protein database [72, 73] were performed for all 63 proteins in the OG, complemented by literature mining using paperBLAST [45], to identify homologues with experimental characterization.

## Declarations

## Availability of data and materials

The datasets used are publicly available and can be found through the original publications cited. All the code used during the current study is available in the git repository: https://github.com/Petra-end/microbial-representations. All supplementary information is also available in this repository, including figures, tables, and model hyperparameters.

## Authors’ contributions

P.M. conceived the study, performed the analyses, visualisation, and wrote the manuscript. G.S. generated the experimental data, gathered the external data, led all bioinformatics analysis and data preprocessing, and manuscript revision. C.M.J.P. and S.A. contributed to project oversight and manuscript revision. R.d.J. supervised the project and contributed to project discussions, interpretation of the results, and manuscript revision. All authors read and approved the final manuscript.

## Acknowledgements

The authors thank dr. Wilson Silva for insightful discussions and informal technical input.

## Ethics approval and consent to participate

Not applicable.

## Consent for publication

Not applicable.

